# TMPRSS2 regulates ACE2 trafficking and shedding

**DOI:** 10.64898/2026.06.17.732908

**Authors:** Yue Qiu, Elena Popova, Oliver Popp, Philipp Mertins, Bernadette Nickl, Fatimunnisa Qadri, Michael Bader

**Author notes:** **Correspondence:** Prof. Dr. Michael Bader, Max-Delbrück-Center for Molecular Medicine (MDC) Robert-Rössle-Str.10, D-13125 Berlin, Germany.

## Abstract

Angiotensin-converting enzyme 2 (ACE2) functions as the receptor for the severe acute respiratory syndrome coronavirus 2 (SARS-CoV-2). The virus utilizes the cellular endocytic machinery for entry by binding to defined residues on ACE2 with its spike protein (S protein), whose activation requires a priming process by another transmembrane protease, the transmembrane protease serine 2 (TMPRSS2). In addition, ACE2 itself is cleaved by TMPRSS2, which has been shown to be critical for viral pathology. This study aimed to elucidate the relationship between ACE2 and TMPRSS2 and the mechanism of ACE2 processing under normal cellular conditions. It is shown that interaction of ACE2 with TMPRSS2 results in altered processing, modification and cellular localization. Glycosylation of ACE2 has a major impact on TMPRSS2 interaction, trafficking and shedding of the enzyme. Studies in newly generated TMPRSS2-knockout rats reveal increased ACE2 levels in tissues supporting an important role of TMPRSS2 in ACE2 shedding also in vivo.

## Introduction

In December 2019, a novel human coronavirus was identified in Wuhan, China^1,2^. The emergence of this SARS-CoV-2 virus has led to a subsequent global outbreak, referred to as the COVID-19 pandemic. Infection with the virus could result in severe respiratory symptoms and lethality, causing millions of confirmed deaths worldwide^3^. The angiotensin-converting enzyme 2 (ACE2) was recognized as the receptor also for the SARS-CoV-2 virus after it had been unveiled as receptor for the SARS-CoV-1 virus in 2003^4–6^.

Full-length ACE2 is a transmembrane protein that functions as a peptidase, transporter, and receptor. ACE2 was initially characterized as a critical mediator of blood pressure regulation by catalyzing the hydrolysis of the vasoconstrictive peptide angiotensin II to the vasodilatory peptide angiotensin-(1-7)^7,8^. During the course of a SARS-CoV-2 infection, the virus infects cells that express ACE2 as a receptor and employs the cellular endocytic machinery to initiate infection. The virus binds to specific residues on ACE2 through the receptor binding domain (RBD) of the spike (S) protein^9^. Processing by Transmembrane protease serine-2 (TMPRSS2) on the surface of the cell results in activation of the S protein and exposure of the fusion peptide of SARS-CoV-1 and SARS-CoV-2, favoring fusion of the viral and host cell membranes and ultimately leading to virus entry^4,5,10^. Notably, ACE2 itself can also be cleaved by TMPRSS2 at its dimerization domain^4,5,10^. The cleavage of both ACE2 and the S protein by TMPRSS2 has been demonstrated to be essential for the process of infection, as only cells that co-express both ACE2 and TMPRSS2 allow for efficient virus entry^5,10,11^.

TMPRSS2 is a member of the type II transmembrane serine protease (TTSP) family, with expression observed in numerous epithelial tissues^12^. It is synthesized as a single-chain zymogen, subsequently undergoing autocatalysis-mediated activation. The mature form of TMPRSS2 is anchored to the cell surface via an N-terminal transmembrane domain^13,14^. It is not only involved in the cellular uptake of SARS-CoV viruses but it also has recently been revealed as direct receptor for the coronavirus HKU1^15^. Moreover, it is involved in the infection of influenza viruses^16,17^ and a recent publication shows that binding of ACE2 may stimulate this process^18^. Accordingly, knockout mice and pigs for TMPRSS2 are protected from infection by influenza and coronaviruses^16,17,19–24^, and the use of the TMPRSS2 inhibitors camostat mesylate and nafamostat as well as newly developed ones to prevent SARS-CoV-2 infection was successful in animal models but still lacks clinical confirmation^25,26^.

Previous research suggests that TMPRSS2 is surface-expressed but undergoes intracellular autoactivation^27^. The authors examined TMPRSS2 zymogen activation and N-glycosylation in biochemical and cellular experiments. However, the cellular mechanism that regulates TMPRSS2 activation and cell surface expression remained elusive.

In this study, we demonstrated that TMPRSS2 can undergo intermolecular transactivation, and that its proteolytic activity is necessary for a productive interaction with ACE2 and for the subsequent processing of ACE2. Activated TMPRSS2 binds to ACE2, inducing two distinct changes in ACE2: a ∼5 kDa mobility shift and cleavage generating an ∼85 kDa N-terminal fragment. However, both effects were absent in the inactive TMPRSS2 mutant (TMPRSS2i). Furthermore, active TMPRSS2 significantly reduced the expression of ACE2 at the cell surface, promoted the accumulation of ACE2 within the intracellular/endoplasmic reticulum (ER)-like compartments, and increased the release of ACE2 into the surrounding medium. Finally, the use of ACE2 N-glycosylation mutants revealed that TMPRSS2 retains the capacity to cleave ACE2 even in the absence of glycan sites. However, the propensity of TMPRSS2-induced membrane trafficking of ACE2 is contingent upon the presence of N3–N7 glycosylation sites. The complete removal of all seven glycosylation sites (N1–N7Q) led to a reduction in ACE2 surface expression and the production of an intracellular pattern that was similar to that of ACE2 co-expressed with TMPRSS2. This suggests that the overall ACE2 glycosylation level is important for ACE2 trafficking and may be linked to how TMPRSS2 alters ACE2 maturation. TMPRSS2 deficient rats exhibit increased tissue levels of ACE2 and reduced expression of inflammation markers in the lung. In conclusion, this study provided substantial evidence to support the hypothesis that activated TMPRSS2 plays a pivotal role in the modulation of ACE2 maturation, trafficking, and shedding.

## Material and Methods

### Generation of TMPRSS2-Knockout rats by CRISPR/Cas9 mediated genome editing

State of Berlin authorities approved the generation of the TMPRSS2-Knockout (KO) rats (crTMPRSS2, license No. G0162/14). The rat model was generated by electroporation of Sprague-Dawley (SD) rat zygotes with a mixture of 1280 ng/µL Cas9 protein (IDT), 258 ng/µL sgRNA with the sequence 5′-GACCUUACUAUGAGAACCACGG (IDT) to target the region in the rat *Tmprss2* gene covering the ATG start codon, and 500 ng/µL tracrRNA (Dharmacon). The electroporated embryos were transferred into foster mothers according to established methods^28^. The offspring were genotyped by polymerase chain reaction with primers flanking the gRNA target region (TMPRSS21, 5′-CCTGTCTTCTGAGTATGACG; TMPRSS22, 5′-TTGTGACTCTCGGAGCATAC) and sequencing of the polymerase chain reaction fragment.

### Plasmids

Full-length hACE2 cDNA with or without GFP tag and TMPRSS2 cDNA with or without FLAG Tag at the C-terminus were cloned into the pcDNA3.1 vector. N-glycosylation mutations of ACE2 were introduced by PCR amplification of synthetic ACE2 cDNA sequence with N103Q, N322Q, N432Q, N546Q, N690Q mutations ordered from GeneArt™. After restriction digestion with HindIII and ClaI, the amplified fragment was cloned into a pcDNA3.1 plasmid already containing ACE2 with and without N53Q and N90Q mutations.

### Cell Culture

HEK293 cells (human embryonic kidney cell) and HeLa cells (human cervical cancer cells) were maintained in DMEM medium (Dulbecco’s modified Eagle’s Medium) containing 10% of FBS (fetal bovine serum) and 1% Penicillin-Streptomycin at 37°C and 5% CO_2_. Passaging of cells was carried out every 3-4 days.

### Expression of recombinant proteins

HEK293 and HeLa cells were transiently transfected with plasmids encoding recombinant proteins using Opti-MEM and Lipofectamine 3000 (Thermo Fisher Scientific). Cell culture medium was collected 24h post transfection and centrifuged at 1500 rpm for 10 min to remove cell pellets. Adherent cells were scraped off the plate and harvested in RIPA buffer (Radio-Immunoprecipitation Assay buffer), transferred to pre-cooled tubes, sonicated for five cycles of 25–30 s at 4°C, and rotated for 30–60 min at 4°C. Lysates were centrifuged at maximum speed for 15 min at 4°C, and the supernatant was collected for further experimental procedures.

### Western Blot

Total protein concentration of cell lysates was measured by BCA (bicinchoninic acid assay. After the concentration was adjusted with PBS buffer, the protein fraction was mixed with sample buffer and separated by SDS-PAGE on 10% acrylamide gels under reducing conditions. Proteins were then transferred onto methanol activated PVDF (polyvinylidene difluoride) membranes (Millipore). After blocking with blocking buffer (LI-COR), the proteins were detected by anti-hACE2 (R&D Systems, AF933, 1:1000), anti-TMPRSS2 (Proteintech, 14437-1-AP, 1:1000), anti-FLAG (Invitrogen, FG4R, 1:500), anti-GAPDH (Invitrogen, PA1-988, 1:1000), anti-ITGβ1 (Santa Cruz Biotechnology, N20,1:500) antibodies. Fluorescence secondary antibodies with IRDye® 800CW or RDye® 680RD (LI-COR, 1:10000) enabled visualization of bands under a LI-COR ODYSSEY® imager.

### Cell membrane protein isolation

Pierce Cell Surface Biotinylation and Isolation kit (Thermo Fisher) was used under manufacturer’s instructions to assess expression of recombinant hACE2 on the surface of HEK293 cells. Cells were cultured in 15cm dishes and transfected with equal amounts of plasmid DNA encoding wildtype or glycosylation-deficient hACE2, and TMPRSS2. After 24 hours incubation, the medium was removed from each dish and cells were washed with 20 mL of PBS. The membrane-impermeable Sulfo-NHS-SS-Biotin was dissolved in PBS. Cells in each dish were biotinylated for 10 min at RT with 10 mL biotin solution, labeling solution was then removed from the dishes and cells were washed and harvested in ice-cold TBS. After centrifugation, 500 μl Lysis Buffer with protease inhibitors was added to the cell pellet. The cells were lysed for 30 min on ice and centrifuged at 15,000 rpm for 5 min. The supernatant was used to isolate labeled proteins by applying 250 µL of it to a NeutrAvidin™ Agarose resin in provided columns and incubated for 30 min at RT on a rotator. The mixture was washed 4 times and then eluted with 250 μl of Elution buffer (10mM DTT). ACE2 surface expression level was evaluated by comparing input and eluate samples by western blot.

### HisPur™ Ni-NTA pull-down of His-tagged proteins

The HEK293 cells were transiently infected with plasmids expressing His-tagged ACE2. Cells were washed once with PBS and lysed on the culture plate using 70–80 µL lysis buffer per well, or 100 µL for highly confluent samples. The lysis buffer did not contain EDTA or reducing agents. Cell lysates were processed as previously described and the cleared supernatant was transferred to a fresh tube. An aliquot of 20–30 µL was reserved as input.

For each sample, 20 µL of HisPur™ Ni-NTA resin was equilibrated with equilibration buffer and pelleted at 700 × g for 2 min. Lysates were mixed with an equal volume of equilibration buffer, added to the resin, and rotated end-over-end for 30 min. The resin was then pelleted at 700 × g for 2 min, and the supernatant was collected as the unbound fraction. The resin was washed thrice with wash buffer and eluted with one resin-bed volume of elution buffer.

SDS sample buffer was added to the collected fractions, and samples were heated at 95°C before SDS–PAGE and Western blot analysis.

### GFP co-immunoprecipitation

The HEK293 cells were transiently infected with plasmids expressing ACE2-GFP fusion proteins and flag-tagged TMPRSS2. Cell pellets were lysed in 200 µL of ice-cold lysis buffer supplemented with protease inhibitors. Cell lysates were processed as previously described and the supernatant was subsequently transferred to a pre-cooled tube and diluted with 300 µL of dilution buffer containing protease inhibitors. (50 µL of this solution was reserved as the input sample). The GFP-binding beads were then gently resuspended, for each sample, 25 µL of the bead slurry was equilibrated in 500 µL of ice-cold dilution buffer. The slurry was then pelleted at 2,500 × g for 5 minutes at 4°C. The diluted lysate was subsequently added to the beads and rotated end-over-end for a period of one hour at a temperature of 4°C. The beads were washed thrice with 500 µL of wash buffer (2,500 × g for 5 min at 4°C after each wash to sediment the beads). The proteins were then eluted by resuspending the beads in 80 µL of 2× SDS sample buffer and boiling them for five minutes at 95°C. Following the pelleting of beads (2,500 × g, 2 min, 4°C), the eluates were subjected to analysis by SDS–PAGE and Western blotting.

### Endoplasmic Reticulum (ER) Staining and Microscopy imaging

Cells were stained with Cell Navigator® Live Cell ER Staining Kit (ER Tracer™ Blue, AAT Bioquest, #22634). A 500× stock was prepared by adding 20 µL DMSO to the ER Tracer™ Blue dye and stored at −20 °C in single-use aliquots. For staining, a working solution was made fresh by diluting the stock 1:500 in Live Cell Staining Buffer (2 µL stock in 1 mL buffer). HeLa Cells in µ-Slide plates (µ-Slide 8 well, Ibidi, #80821) were incubated with the working solution (50 µL per well, respectively) at 37 °C for 30 min protected from light. The solution was removed, cells were washed three times with PBS, and fluorescence was imaged using a microscope (Leica TCS SP8 WLL) with GFP DAPI filter. If needed, cells were fixed after staining with 4% formaldehyde for 10 min and washed three times before imaging.

### Protein isolation from organs

The excision of the organs was followed by immediate freezing in −40°C cold isopentane on dry ice until processing. For homogenization, pre-labeled bead-containing tubes (5 beads per tube, added with forceps) were prepared and kept on ice. All steps were carried out on ice with pre-chilled buffers. Each organ sample was placed into 500 µL of the appropriate buffer depending on the downstream application: MES assay buffer for ACE2 activity measurements (50 mM MES, 300 mM NaCl, 10 µM ZnCl2, 1.2 µM HCl, 0.01% Brij-35, pH 6.5), homogenization buffer for TMPRSS2 activity measurements (25 mM sodium phosphate, pH 6.8, 2 mM DTT, 0.5 mM EDTA), or RIPA buffer supplemented with EDTA for immunoblotting. Tissues were mechanically homogenized twice for 40 s and returned to ice immediately after each cycle. Subsequent to these cycles, the samples underwent sonication which consisted of five cycles of 30 seconds at a high-power setting. Lysates were cleared by centrifugation at maximum speed for 10 min at 4°C. The resulting supernatants were collected for further analyses.

### Measurement of ACE2 Activity

The activity of the ACE2 enzyme was measured using a fluorogenic peptide substrate in a 96-well plate format. Reactions (100 µL per well, performed in duplicate) were prepared in MES assay buffer in black, flat-bottom 96-well plates. The fluorogenic substrate MCA–YVADAPK was supplied as a 400 µM stock solution in DMSO and added at 2 µL per reaction to yield a final concentration of 8 µM. In instances where indicated, the ACE2 inhibitor MLN-4760 was added to a final concentration of 100 nM and samples were pre-incubated with the inhibitor in assay buffer for 15 min at ambient temperature prior to substrate addition. Fluorescence was monitored at 37°C using a microplate reader (INFINITE M PLEX, TECAN Germany) with excitation at 320 nm and emission at 405 nm.

### Measurement of TMPRSS2 Activity

TMPRSS2 enzymatic activity was measured using a fluorogenic AMC-based peptide substrate in a 96-well plate format. Reactions (100 µL per well, performed in duplicate) were assembled in black, flat-bottom 96-well plates using 20 µL of 5× assay buffer (0.1 M Tris-HCl, pH 8.2, 0.1 M CaCl2) and brought to final volume with the sample solution and water as needed. The fluorogenic substrate, Boc-Gln-Ala-Arg-7-aminomethylcoumarine (AMC)·HCl was prepared as a 5 mM stock solution (50×) in water and diluted to 5× working strength immediately prior to use. A volume of 20 µL of the 5× substrate solution was added to each well to initiate the reaction. In instances where indicated, the TMPRSS2 inhibitor nafamostat was added at a final concentration of 5 µM. Fluorescence was monitored at 37°C using TECAN microplate reader (INFINITE M PLEX, TECAN Germany) with excitation at 380 nm and emission at 440 nm.

### Proteomics methods

#### Proteomic sample preparation and LC–MS/MS analysis

For proteomic analysis, lung tissue protein extracts from wild-type (WT) and crTMPRSS2 rats were analysed at the Proteomics Technology Platform of the Max Delbrück Center (MDC, Berlin, Germany). Approximately 2–4 mg of cryopulverised tissue was collected from the CryoPrep bag and lysed in 1% (w/v) sodium deoxycholate (SDC), 100 mM Tris-HCl (pH 8.5), 150 mM NaCl, 1 mM EDTA, 10 mM dithiothreitol (DTT), and 40 mM chloroacetamide (CAA). Lysis, reduction, and alkylation were performed simultaneously by heating at 90–95°C for 10 min. Proteins were digested overnight at 37°C using Lys-C and trypsin (1:50 enzyme-to-protein ratio). Digestion was stopped by acidification with formic acid (FA; final concentration 1%), precipitating SDC. Following centrifugation, peptides were desalted on an AssayMAP Bravo automated platform (Agilent Technologies), dried, and reconstituted in 3% acetonitrile (ACN)/0.1% FA.

Peptides were separated on a Vanquish Neo UHPLC system (Thermo Fisher Scientific) coupled online to an Orbitrap Astral mass spectrometer (Thermo Fisher Scientific) operated in positive-ion nano-electrospray mode. Chromatographic separation was performed at 0.35 μL/min using solvent A (3% ACN, 0.1% FA) and solvent B (80% ACN, 0.1% FA) with a 26 min gradient: 2% B at injection, 7% B at 1 min, 20% B at 11 min, 30% B at 18.5 min, and 60% B at 20.5 min, followed by a wash at 90% B from 21.5–26 min and column re-equilibration. DIA data were acquired using an Orbitrap MS1 survey scan (m/z 380–1100, resolution 240,000, AGC target 500%, 3 ms injection time) followed by Astral DIA scans covering precursor m/z 380–980 with 2 m/z isolation windows, HCD fragmentation at 25% normalised collision energy, a scan range of m/z 150–2000 and 3 ms injection time, with a cycle time of 0.6 ms.

#### DIA-NN data processing

Raw DIA data were processed using DIA-NN^29^ with a predicted spectral library generated from the UniProt *Rattus norvegicus* isoform reference proteome (UP000002494, 202404) supplemented with a universal protein contaminants database. In silico digestion was performed with tryptic specificity after lysine and arginine, allowing up to two missed cleavages and peptide lengths of 7–30 amino acids. Carbamidomethylation of cysteine (UniMod:4) was specified as a fixed modification, while methionine oxidation (UniMod:35) and protein N-terminal acetylation (UniMod:1) were allowed as variable modifications (maximum one variable modification per peptide). Methionine excision, peptidoform inference, match-between-runs, and retention-time profiling were enabled. Identification confidence was controlled at 1% false discovery rate (FDR).

#### Differential protein abundance analysis

Downstream statistical analysis was performed in R. Potential contaminant proteins were removed and protein group intensities were log₂-transformed. Protein groups supported by fewer than two identified peptides were excluded prior to analysis. Sample relationships and data quality were assessed by principal component analysis (PCA).

To retain proteins with sufficient quantitative information, valid-value filtering was applied such that a protein was retained if at least three valid (non-missing) measurements were present in at least one experimental group (OR logic; minimum 3 biological replicates per group). Missing values were subsequently imputed using a left-shifted intensity-based imputation approach for visualisation and statistical testing.

Differential abundance analysis between WT and crTMPRSS2 groups was performed using the limma package^30^. For each protein, a linear model was fitted to the log₂-transformed intensity values, and moderated t-statistics were calculated using empirical Bayes variance moderation with intensity-dependent trend correction and robust variance estimation (trend = TRUE, robust = TRUE). P-values were adjusted for multiple testing using the Benjamini–Hochberg procedure, and proteins with an adjusted p-value < 0.05 were considered significantly differentially abundant.

#### Peptide coverage analysis

Distinct peptide quantification was performed using the DIA-NN report.parquet output. Precursor entries were retained only when Q.Value, Protein.Q.Value, Lib.Q.Value, and Lib.PG.Q.Value were all ≤ 0.01. For each peptide sequence within a run, the precursor exhibiting the highest Precursor.Normalised intensity was selected (Top1 approach) to obtain a single quantitative value per peptide and sample. No additional normalisation was applied during downstream analysis.

Residue-resolved peptide coverage maps were generated with R. For each identified peptide, the measured intensity was divided by peptide length to obtain a per-residue contribution. Peptides were mapped to the corresponding protein sequence by exact string matching, including repeated sequence occurrences, and per-residue intensities were summed across all overlapping peptides to generate quantitative coverage profiles along the protein sequence.

### RNASeq

For transcriptomic profiling, total RNA was extracted from lung tissues of WT and crTMPRSS2 rats and submitted to Novogene (Munich, Germany) for RNA sequencing. RNA integrity assessment, library preparation, sequencing, and initial data processing were performed by the service provider according to their standard workflow.

### Functional enrichment analysis

GO Biological Process enrichment analysis was performed to identify biological functions associated with transcriptional changes in crTMPRSS2 rat lung tissue. Differentially expressed genes from the RNA-seq comparison between crTMPRSS2 and WT rats were clustered using an adjusted p-value ≤ 0.05 and |log2FC| ≥ 1. Up-and down-regulated genes were analyzed separately.

### RT-qPCR

RT-qPCR was used to assess gene expression in cells and tissues. Reactions were executed in 384-well plates on a QuantStudio 5 instrument with GoTaq® qPCR Master Mix, SYBR Green® for fluorescence detection, and CXR as a passive reference dye. Primers were designed to yield amplicons with a size of less than 200 base pairs (bp) and comparable melting temperatures. This design enables the amplification of DNA at a uniform annealing temperature of 60°C. Each reaction contained 9 ng cDNA, and samples were measured in duplicate. The specificity of the assay was confirmed by melting-curve analysis, and reactions that produced non-specific products or primer-dimers were excluded from further analysis. The subsequent data processing was conducted using QuantStudio™ Design & Analysis Software v1.3.1, with the threshold settings being manually configured during the exponential phase. The expression levels were then normalized to Tbp, Hprt, or 18S rRNA and calculated using the 2^-ΔCt method.

### Immunofluorescence

For the immunofluorescence staining 5µm thick lung paraffin sections were deparaffinized, rehydrated and boiled in antigen retrieval buffer either in Tris/EDTA pH 9.0 (Lcn2) or Citrate buffer pH 6.0 (SFTPC, CD68) for 15 min. Nonspecific bindings were blocked using a mixture of 10% normal donkey serum and 2% bovine serum albumin at room temperature for 30 min. For the immune staining the sections were incubated with following antibodies; anti-SFTPC (1:500, Cat. #ABC99, Merck), anti-CD68 (1:500, Cat. #MCA341R, Serotec) and anti-Lcn2 (1:1000, Cat. #Ab216462, Abcam) over night at 4°C in a wet chamber. Next day the sections were washed and incubated with fluoresceine-conjugated secondary antibodies in a concentration of 1:300 for 2 h at RT, washed and cover-slipped with Fluoromount-G with DAPI mounting medium (Cat. #00-4959-52, Invitrogen). The sections were examined with an inverted bright light/fluorescence microscope (Keyence BZ-X800, Germany).

### Statistics

All data are expressed as mean ± standard deviation (SD). Statistical differences between two groups were assessed using an unpaired, two-tailed Student’s t-test. Differences were considered statistically significant at p < 0.05. Statistical analyses were carried out using Python (SciPy package).

## Results

### TMPRSS2 autoactivation happens in trans

TMPRSS2 exerts proteolytic activity only after maturation, which is an autoactivation process that cuts the protease zymogen into 2 fragments, and the enzymatically reactive serine protease domain (SPD) remains connected to the stem region through a disulfide bond after cleavage. We wanted to explore whether this autoactivation happens intramolecular (in cis) or intermolecular (in trans). Co-transfections of TMPRSS2 with a proteolytically inactive TMPRSS2 mutant (TMPRSS2i) showed that this autoactivation could happen in trans, since TMPRSS2i-FLAG can also be cleaved by active TMPRSS2 without tag, and the released SPD signal can be recognized with a FLAG antibody (Figure 1). This confirms a recent study and suggests dimerization of TMPRSS2, which could be relevant to its function^31^.

**Figure 1.**
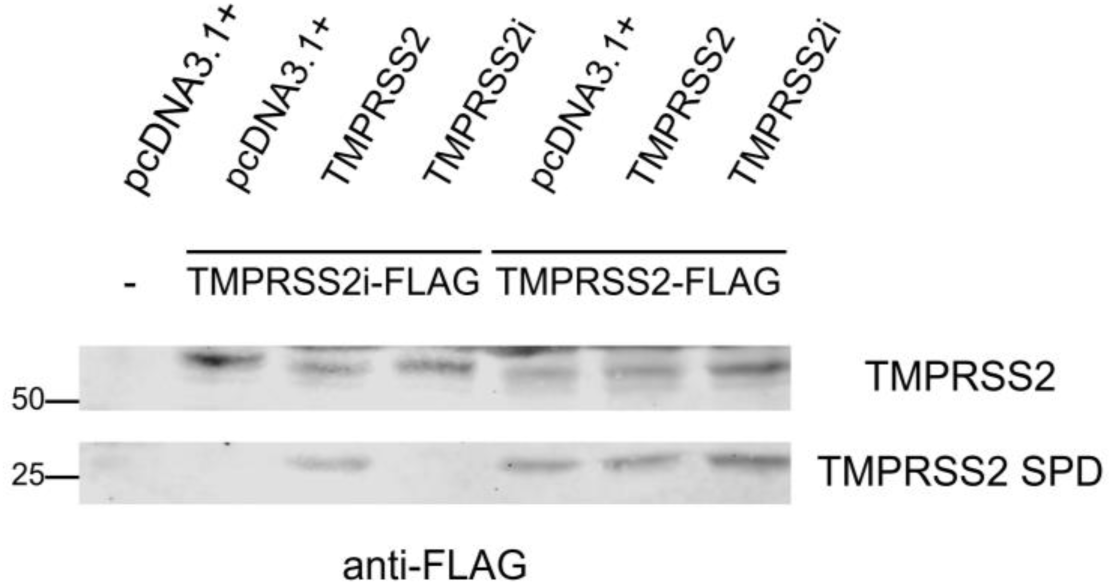
TMPRSS2 undergoes intermolecular trans-autoactivation. To determine whether TMPRSS2 maturation occurs in cis or in trans, cells were co-transfected with active, untagged TMPRSS2 and catalytically inactive FLAG-tagged TMPRSS2 (TMPRSS2i-FLAG).

### Activated TMPRSS2 binds to ACE2 and cleaves and alters ACE2 processing

In order to study the interaction of ACE2 and TMPRSS2, HEK293 cells were transiently transfected with plasmids expressing ACE2 and co-transfected with TMPRSS2 or TMPRSS2 mutants including TMPRSS2i, N-terminal 1-83 amino acids deleted TMPRSS2 (del1-83TMPRSS2) and N-terminal 1-83 amino acids deleted inactive TMPRSS2 (del1-83TMPRSS2i). The N-terminal 83 amino acids constitute the cytoplasmic domain of the enzyme^32^ and we interrogated its role in the interaction with ACE2.

Co-expression of ACE2 with TMPRSS2 after 24 hours resulted in a shift in the size of ACE2 of approximately 5 kDa on the SDS PAGE, and a cleavage of ACE2 producing a roughly 85 kDa N-terminal fragment. Both the shift and cleavage of ACE2 depended on the proteolytic activity of TMPRSS2, as neither of the two effects were observed when ACE2 was co-transfected with the enzymatically inactive mutant, TMPRSS2i. The N-terminus deleted version del1-83TMPRSS2 was able to induce the shift but only less efficiently the cleavage, suggesting that these two ACE2 changes may arise from separable mechanisms (Figure 2a).

**Figure 2.**
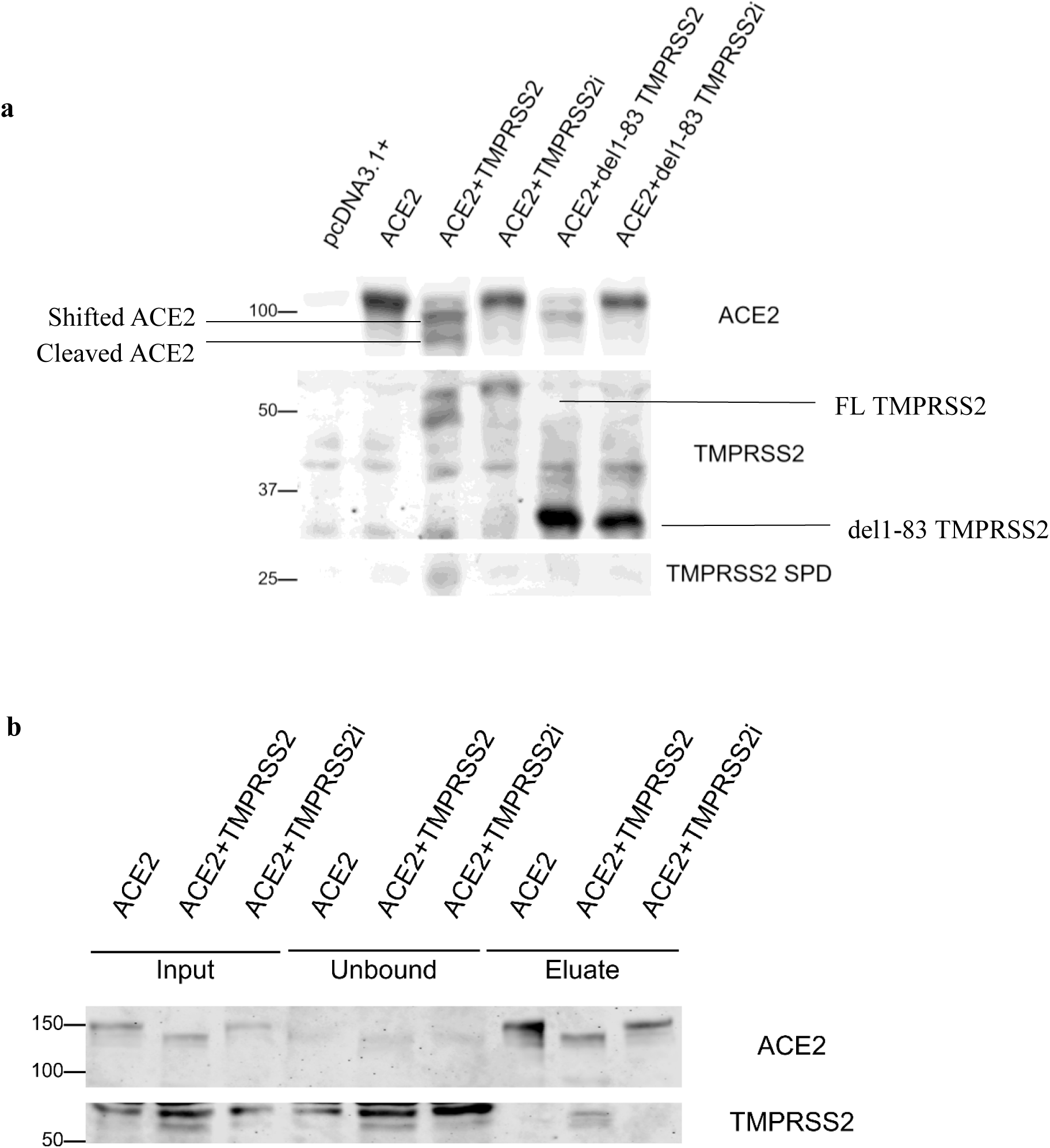
TMPRSS2 activity drives ACE2 processing and ACE2 association. **(a)** HEK293 cells were co-transfected with ACE2 and WT TMPRSS2 or the indicated mutants. After 24 h, WT TMPRSS2 caused an ∼5 kDa shift in ACE2 and generated an ∼85 kDa N-terminal ACE2 fragment. The inactive mutant (TMPRSS2i) did not, and del1–83TMPRSS2 produced the shift but not the cleavage. **(b)** HisPur™ Ni-NTA pull-down of His-tagged ACE2 shows co-precipitation of TMPRSS2 only in its active form, while TMPRSS2i does not co-precipitate.

Using HisPur™ Ni-NTA resin, immunoprecipitation of the His-tagged ACE2 protein revealed that TMPRSS2 only co-precipitated with ACE2 in its active form. However, TMPRSS2i was unable to form a tight interaction with ACE2 (Figure 2b). Taken together, these results indicate that TMPRSS2 association with ACE2 is contingent on TMPRSS2 protease activation: only catalytically active TMPRSS2 co-precipitated with ACE2 and concomitantly induced both the ∼5 kDa mobility shift and ACE2 cleavage. Thus, activated TMPRSS2 binds ACE2 and, through its proteolytic activity, cleaves ACE2 and remodels ACE2 processing.

### ACE2 membrane compartmentalization is reduced by TMPRRS2

ACE2 functions as a peptidase and receptor on the cell membrane, and we were interested how the presence of TMPRSS2 affects ACE2 membrane compartmentalization. ACE2 fused to green fluorescent protein (GFP) was co-transfected with TMPRSS2 and the different mutants in HeLa and HEK293 cells. 24 hours after, the transfected HeLa cells were labelled with a fluorescent kit staining ER and confocal microscopy imaging was performed (Figure 3a). Surprisingly, ACE2 expressed alone showed distinct cell membrane localization, while co-expression with TMPRSS2 resulted in reduced membrane localization and retention of ACE2 in the cytoplasm. Transfection of TMPRSS2i did not induce this retention, but the overall expression of ACE2 on the membrane was also slightly reduced. Since we saw strong correlation of the GFP localization and blue ER fluorescence in Hela cells, we assume that ACE2 was retained in the ER for processing by TMPRSS2. However, as soon as it reached the plasma membrane it was released by TMPRSS2 shedding. The interaction with ACE2 is dependent on the enzymatic activity of TMPRSS2. To further verify this hypothesis, 24 hours transfected HEK293 cells were biotinylated on the membrane using a commercially available kit, the membrane proteins were pulled down and loaded on western blots. Consistent with our microscopic observation, membrane ACE2 was only clearly detectable when transfected alone or together with mutant TMPRSS2i. The ACE2 membrane expression was largely depleted in the presence of active TMPRSS2 (Figure 3b). Accordingly, release of ACE2 in the medium was detected when co-expressed with TMPRSS2 and less with del1-83TMPRSS2 (Figure 3d). No cytoplasmic contamination was seen as the GAPDH signal decreased along washing steps and was not detectable in the eluate (Figure 3a, c), and total amount of cell membrane protein was shown to be consistent after pull-down by an equal ITGβ1 signal in all conditions.

**Figure 3.**
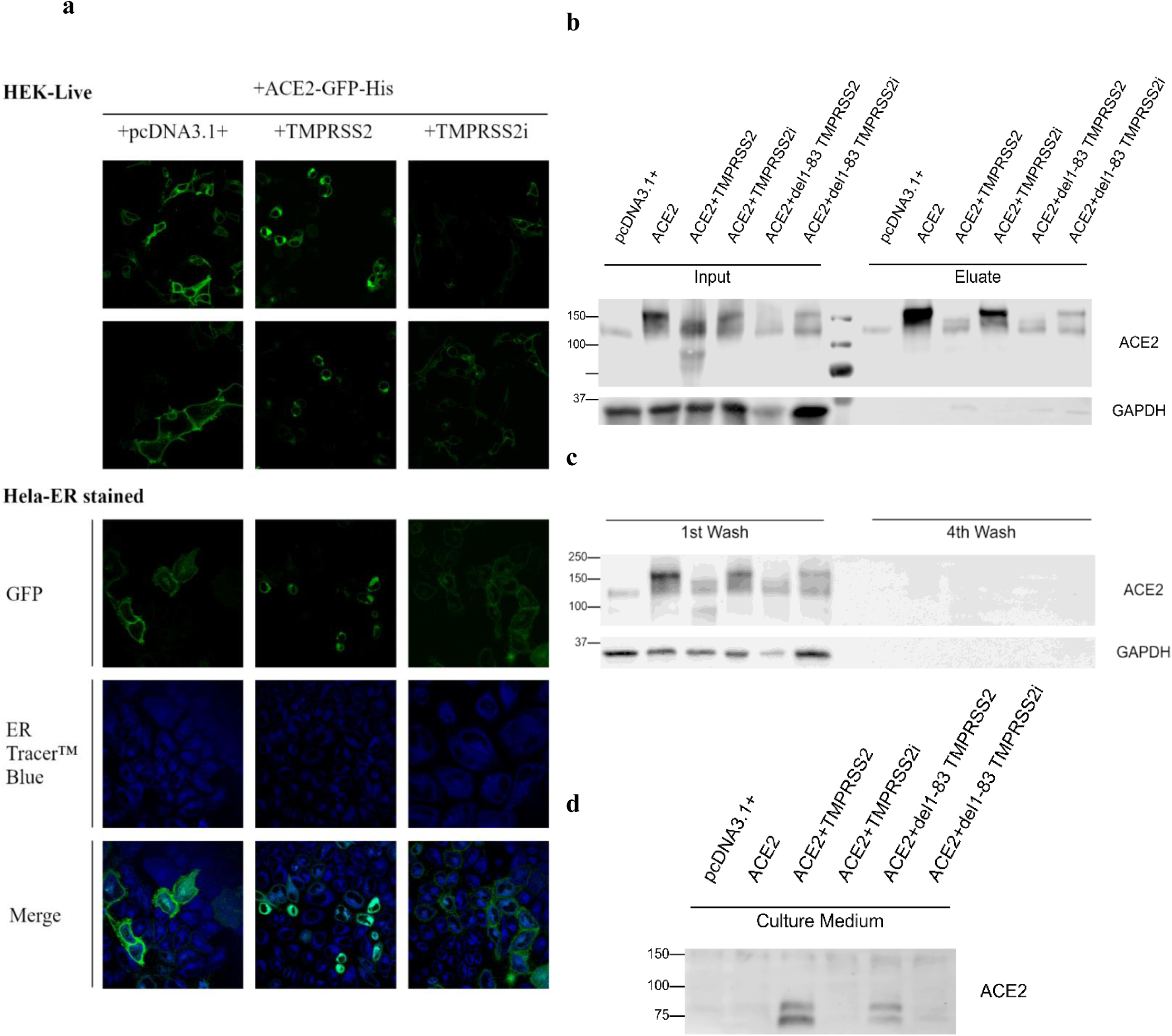
Active TMPRSS2 depletes ACE2 from the cell surface and increases ACE2 release. **(a)** Confocal imaging of HeLa cells 24 h after transfection with ACE2-GFP ± TMPRSS2 (WT or mutant) and ER staining. ACE2-GFP alone localized mainly at the plasma membrane, while co-expression with active TMPRSS2 shifted ACE2-GFP to a cytoplasmic/ER-like pattern. **(b)** Surface biotinylation and pull-down of membrane proteins from HEK293 cells show that surface ACE2 is reduced with active full-length and del1-83 TMPRSS2, but remains detectable when ACE2 is expressed alone or with TMPRSS2i. **(c)** GAPDH control signal disappeared along washing steps indicating no cytoplasmic contamination in the membrane fraction/eluate. **(d)** ACE2 is detected in the medium upon co-expression with active full-length and del1-83 TMPRSS2, consistent with ACE2 shedding.

### Effect of N-glycosylation sites on the interaction between ACE2 and TMPRSS2

ACE2 has 7 N-linked glycosylation sites. With average mass of a single glycan of around 4 kDa^33^, the total glycan mass on ACE2 amounts to about 30 kDa^34^. The majority of glycans at N53, N90, N103, N322, N432, N546, and N690 residues of ACE2 are of the complex-type. The glycans occupy more than 75% of the residues, and the sialic acid linkage always exists in the glycans^35^. As glycosylation have been reported to play an important role in the binding of ACE2 with the SARS-CoV-2 spike protein^36^, we attempted to investigate whether it could also affect ACE2 interaction with TMPRSS2. To this purpose, three ACE2 glycan deficient mutants have been cloned, lacking the N53 and N90 (N1-N2Q), the N103, N322, N432, N546, and N690 (N3-N7Q), and all 7 glycosylation sites (N1-N7Q). All three mutants were efficiently expressed and migrated to the expected molecular weight. Although all three mutants can be effectively cleaved by TMPRSS2, the shift in the apparent size of WT ACE2 induced by TMPRSS2 was no longer observed in ACE2 N3-N7Q and N1-N7Q (Figure 4a), suggesting that this shift was due to reduced glycosylation on N3-N7 caused by structural hindrance upon TMPRSS2 binding.

**Figure 4.**
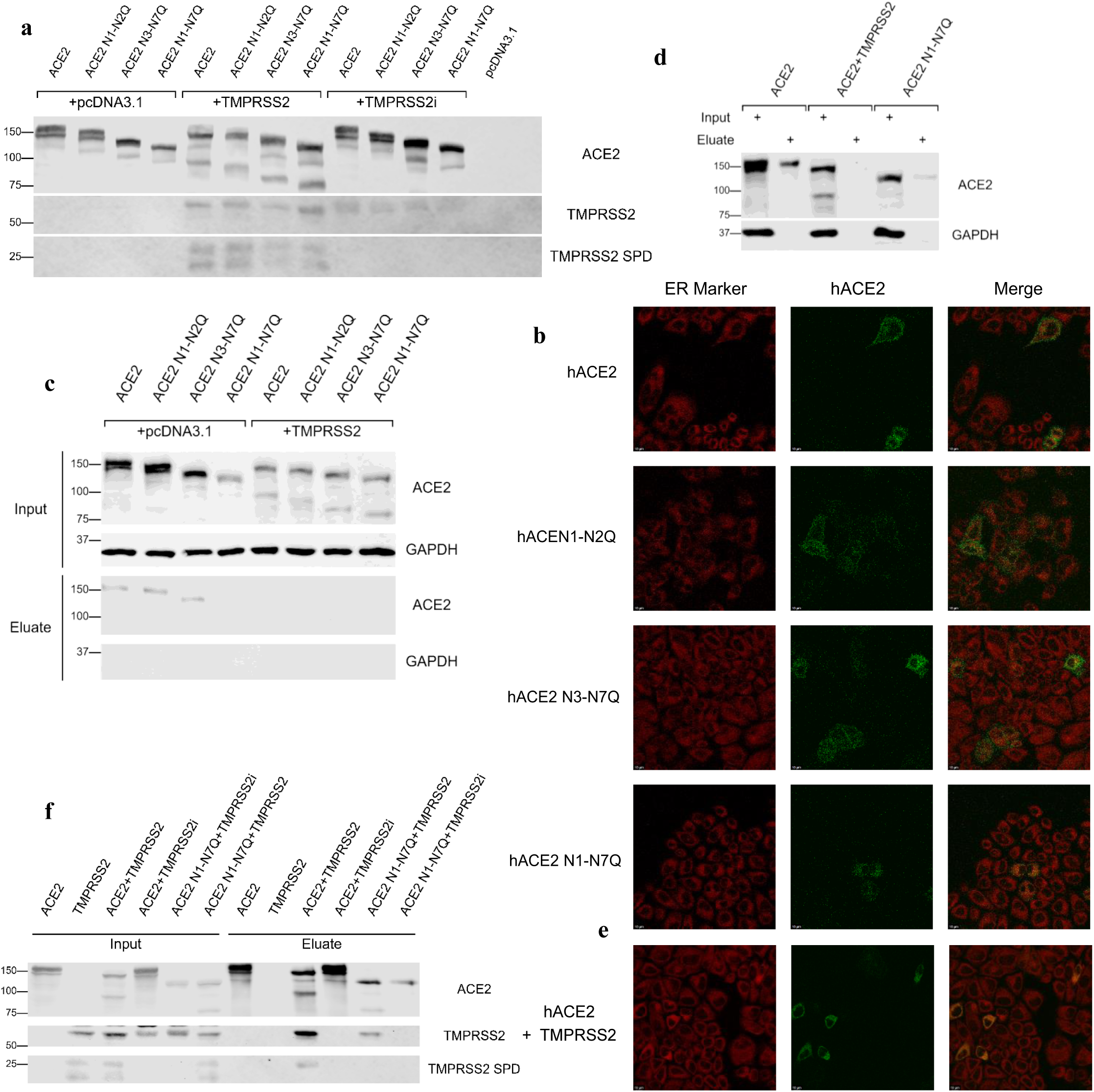
ACE2 N-glycosylation controls the TMPRSS2-induced mobility shift and ACE2 surface localization. **(a)** WT ACE2 and glycosylation mutants (N1–N2Q, N3–N7Q, N1–N7Q) were expressed ± TMPRSS2 (WT or mutant). All mutants were cleaved by TMPRSS2, but the TMPRSS2-induced ACE2 mobility shift was lost in N3–N7Q and N1–N7Q. **(b)** Confocal imaging of ACE2–GFP in HeLa and HEK293 cells. WT ACE2 localized at the plasma membrane, while N1–N7Q showed reduced surface localization; N1–N2Q and N3–N7Q remained largely surface-expressed. **(c)** Surface biotinylation confirms reduced membrane ACE2 for N1–N7Q, but not for N1–N2Q or N3–N7Q. Coexpression with TMPRSS2 diminished ACE2 membrane expression in all cases. **(d–e)** ACE2 N1–N7Q shows an intracellular pattern similar to WT ACE2 co-expressed with TMPRSS2 with ER retention. **(f)** Immunoprecipitation of GFP-tagged ACE2 revealed markedly reduced TMPRSS2 binding to glycan-deficient ACE2 and loss of co-precipitation of the TMPRSS2 SPD domain.

### Effect of N-glycosylation sites on membrane compartmentalization of ACE2

Next we wanted to assess the extent to which glycosylation influences the membrane compartmentalization of ACE2. To this end, the three ACE2 glycosylation mutants were fused with a GFP tag, and their subcellular localization was imaged using a fluorescence confocal microscope in both fixed HeLa cells and live HEK293 cells. Wildtype ACE2 exhibited distinct cell surface localization, while the N1-N7Q demonstrated a significant decrease in membrane expression. However, neither ACE2 N1-N2Q nor ACE2 N3-N7Q was depleted on the cell surface, indicating that the total amount of glycosylation, rather than specific mutations, might be the determining factor of ACE2 surface trafficking (Figure 4b). Membrane protein isolation using cell-surface biotinylation yielded consistent results. A significant reduction in ACE2 membrane localization was observed in ACE2 N1-N7Q, while no such reduction was evident in N1-N2Q and ACE2 N3-N7Q (Figure 4c). The expression pattern of N1-N7Q exhibited a high degree of similarity to that of the WT ACE2 when coexpressed with TMPRSS2 both in membrane protein isolation and microscopic imaging (Figure 4d, e), suggesting retention of ACE2 in the ER.

To further examine the influence of ACE2 glycosylation on its interaction with TMPRSS2, immunoprecipitation was used to analyze the co-precipitation of TMPRSS2 in the presence of glycan-deficient ACE2 mutants. In comparison with the WT ACE2, the non-glycosylated ACE2 N1-N7Q variant exhibited a significantly diminished association with TMPRSS2. It is noteworthy that the TMPRSS2 serine protease domain (SPD), which was readily detected in complexes with WT ACE2, was no longer observed to co-precipitate with the glycan-deficient mutant (Figure 4f). These findings suggest that the loss of ACE2 glycosylation substantially weakens TMPRSS2 binding and indicate that N-linked glycans on ACE2 contribute to stabilizing or facilitating the interaction between the two proteins. Collectively, these results lend support to the hypothesis that ACE2 glycosylation plays a significant role in the formation of the TMPRSS2–ACE2 complex.

### Loss of TMPRSS2 is associated with ACE2 tissue retention

Our studies in cell culture suggested TMPRSS2 to be a sheddase for ACE2 releasing it from cells and tissues. In order evaluate the *in-vivo* relevance of these findings, we generated a TMPRSS2 KO rat model by CRISPR/Cas9 technology carrying a 2A insertion predicted to cause a frameshift in exon 2 of the *Tmprss2* transcript (Figure 5). In these crTMPRSS2 rats, the activity of TMPRSS2 was measured using a fluorogenic peptide substrate for TTSP enzymes, Boc-Gln-Ala-Arg-AMC·HCl, revealing reduced activity in lung and bladder tissue in comparison with WT controls (Figure 6a, b). The activity was not abolished, since other TTSP family enzymes are still present in these rats and are also inhibited by nafamostat. In order to prove the absence of TMPRSS2, we performed proteomics experiments. Due to the low expression of TMPRSS2 in the lung, proteomics discovered only one specific peptide in all wildtype rats which was absent in all crTMPRSS2 animals (Figure 7).

**Figure 5.**
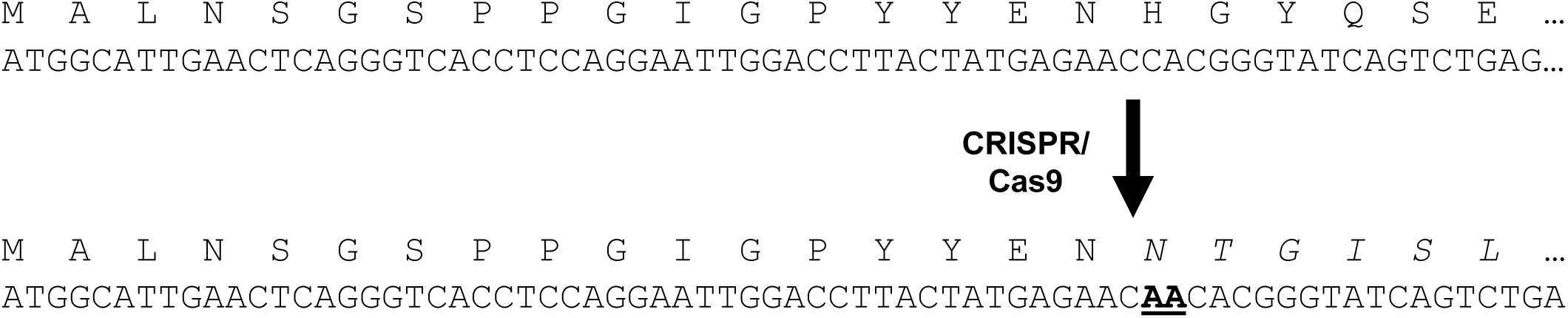
Generation of the crTMPRSS2 rat model. A TMPRSS2 knockout rat was generated by CRISPR/Cas9-mediated genome editing. The mutant allele carries a 2A insertion in the *Tmprss2* locus, which is predicted to cause a frameshift and disrupt normal TMPRSS2 expression and function.

**Figure 6.**
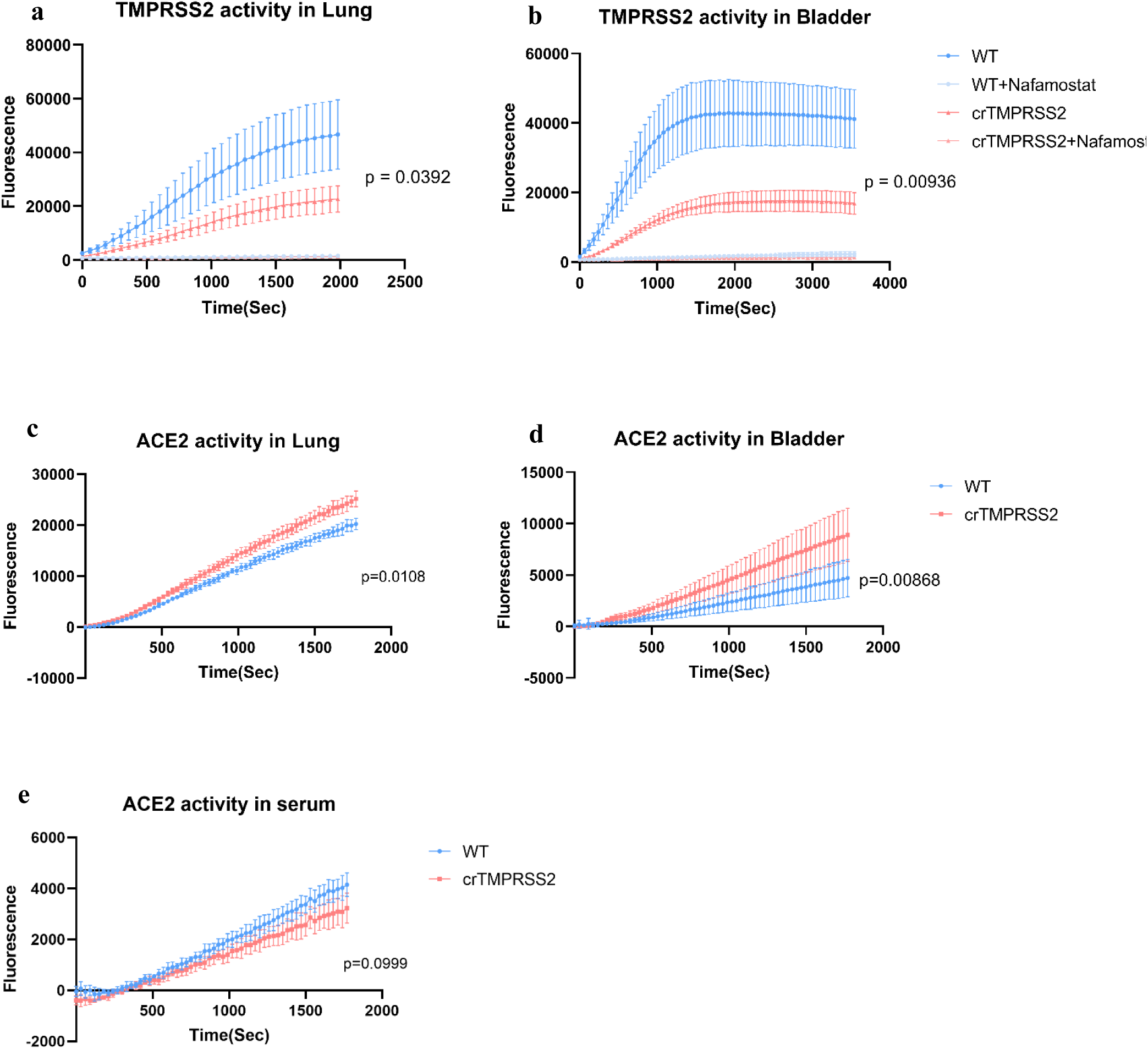
Loss of TMPRSS2 is associated with reduced TMPRSS2-like protease activity and increased tissue ACE2 activity. **(a, b)** Protease activity in lung (a) and bladder (b) lysates from wild-type (WT) and crTMPRSS2 rats was measured using the fluorogenic substrate Boc-Gln-Ala-Arg-AMC·HCl. Substrate cleavage was significantly reduced in crTMPRSS2 animals compared with WT controls in both lung (*p* = 0.0392) and bladder (*p* = 0.00936), consistent with depleted TMPRSS2 activity. Residual activity remained detectable, likely due to other related proteases. **(c-e)** ACE2 activity was measured using the fluorogenic substrate MCA–YVADAPK in lung (c), bladder (d), and serum (e). ACE2 activity was significantly increased in lung (*p* = 0.0108) and bladder (*p* = 0.00868) of crTMPRSS2 rats, while serum ACE2 activity showed a non-significant decrease (*p* = 0.0999).

**Figure 7.**
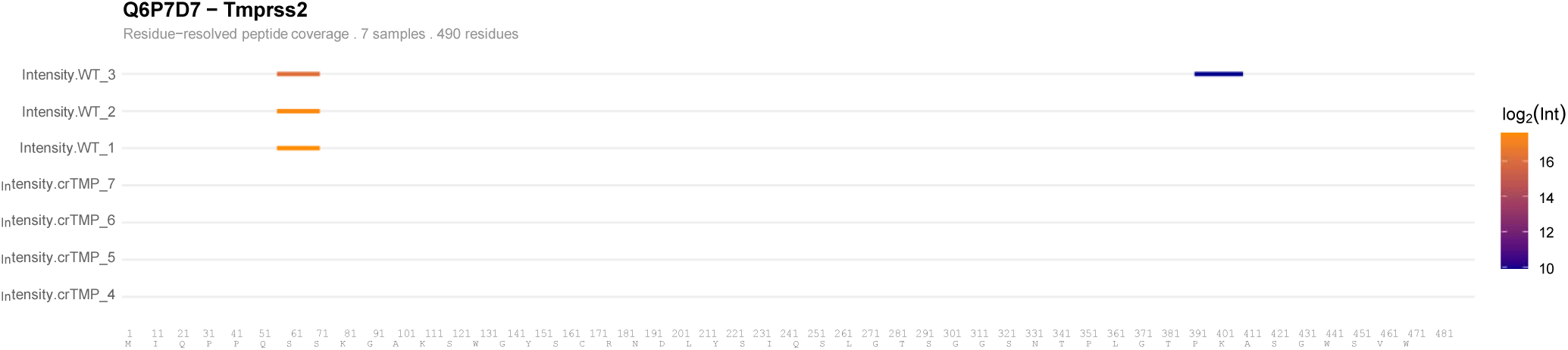
Proteomic detection of TMPRSS2 in WT and crTMPRSS2 lung tissue. Residue-resolved peptide coverage map of *Tmprss2* across individual samples from crTMPRSS2 and WT rats. Each row represents one sample; colour indicates log₂-transformed, length-normalised peptide intensity (dark blue, low; dark orange, high). Positions with no detected peptides are left blank.

The activity of ACE2 was measured using fluorogenic substrate MCA–YVADAPK in lung and bladder tissue and serum of crTMPRSS2 rats and controls. ACE2 activity levels in tissues exhibited a significant increase, while serum levels demonstrated a slight decrease (Figure 6c-e). Western blot analysis showing elevated ACE2 band intensity in bladder tissue from crTMPRSS2 rats further supported this increase in bladder ACE2 at the protein level. (Figure 8). These findings suggest that TMPRSS2 contributes to the shedding and release of ACE2 from tissue into the circulation *in vivo*.

**Figure 8:**
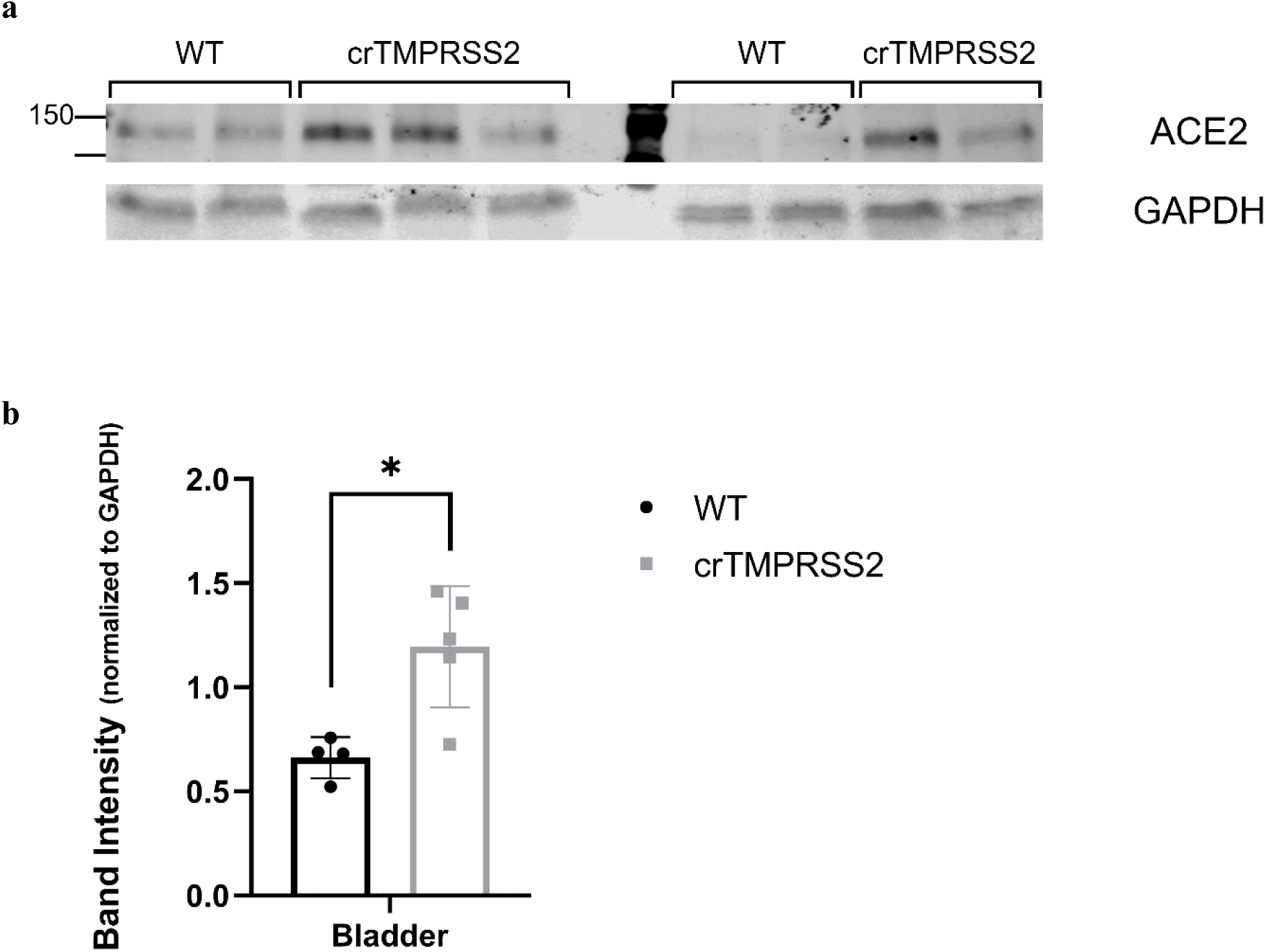
Increased ACE2 protein abundance in bladder tissue of crTMPRSS2 rats. **(a)** Representative Western blot of ACE2 in bladder tissue from WT and crTMPRSS2 rats. GAPDH was used as loading control. **(b)** Quantification of ACE2 band intensity normalized to GAPDH. ACE2 protein levels were significantly increased in bladder tissue of crTMPRSS2 rats compared with WT controls (*p* = 0.0107), consistent with the elevated ACE2 activity detected in this tissue.

### Loss of TMPRSS2 is associated with reduced inflammatory marker expression in the lung

Our proteomics (Figure 9) and RNAseq data (Figure 10) followed by gene ontology analysis (Figure 11) revealed a significant reduction in the expression of inflammatory and innate immune response marker proteins, such as Lcn2 (NGAL) and complement factor C5, and an upregulation of adaptive immune response genes. Both were confirmed by RT-qPCR measurement of mRNA levels (Figure 12). Furthermore, immunofluorescence staining of lung sections revealed a reduction in inflammatory marker expression in crTMPRSS2 lungs compared with WT controls. Lcn2 staining was clearly detectable in WT lung tissue but was markedly reduced or largely absent in crTMPRSS2 sections. The presence of CD68-positive cells was observed in both genotypes; however, the CD68 signal appeared reduced in the alveolar regions of crTMPRSS2 lungs, suggesting decreased macrophage-associated inflammatory activity. SFTPC staining was observed in both WT and crTMPRSS2 lungs, suggesting the maintenance of alveolar type II epithelial cell marker expression. Collectively, these observations indicate that the loss of TMPRSS2 is associated with diminished pulmonary inflammatory marker expression, particularly as evidenced by reduced Lcn2 and alveolar CD68 staining (Figure 13).

**Figure 9.**
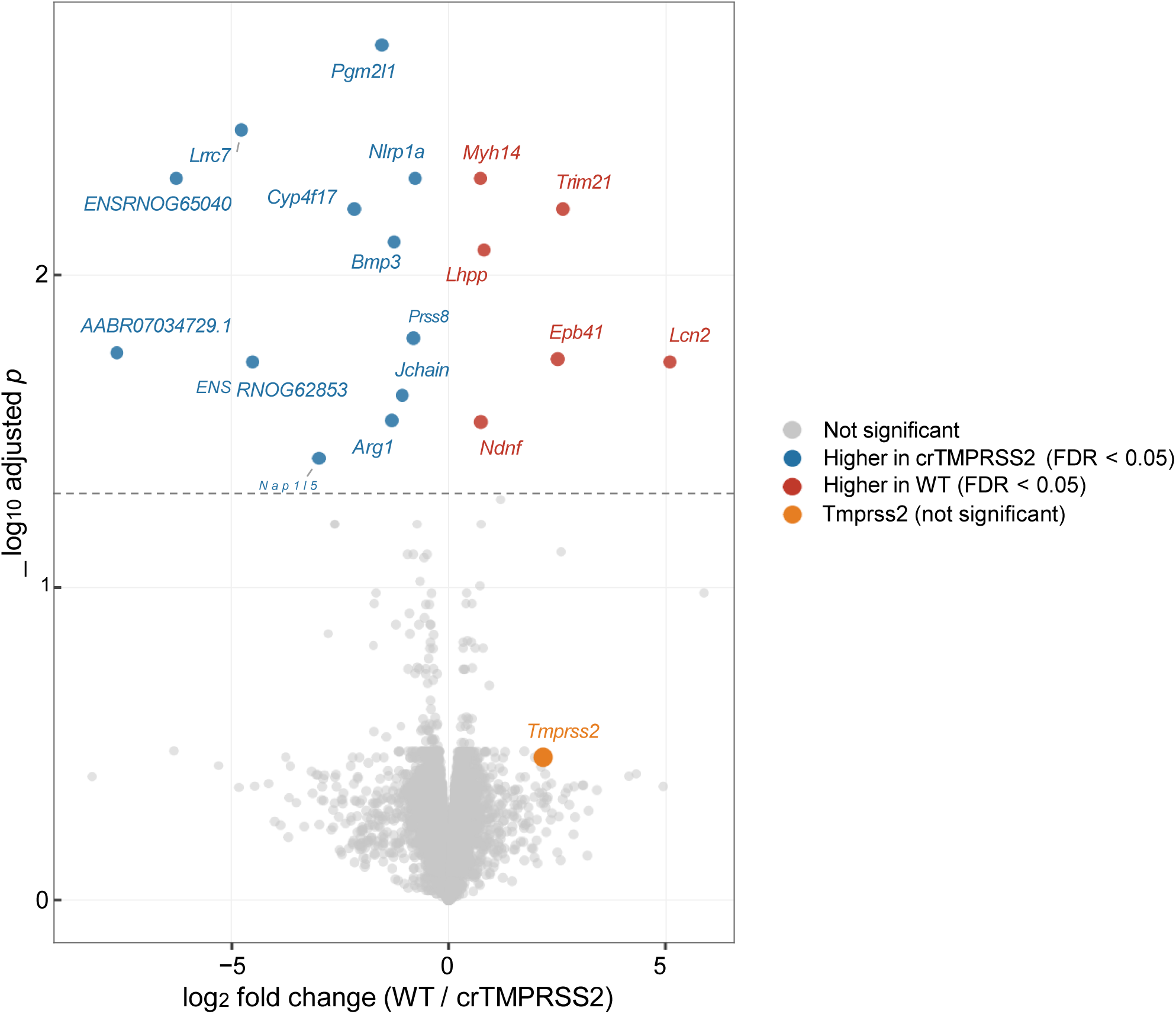
Proteomic analysis of lung tissue from crTMPRSS2 and WT rats. Volcano plot showing differential protein abundance in lung tissue of crTMPRSS2 rats compared with wild-type (WT) controls. Significantly altered proteins are highlighted according to the indicated thresholds. Lcn2 was reduced in crTMPRSS2 lungs.

**Figure 10.**
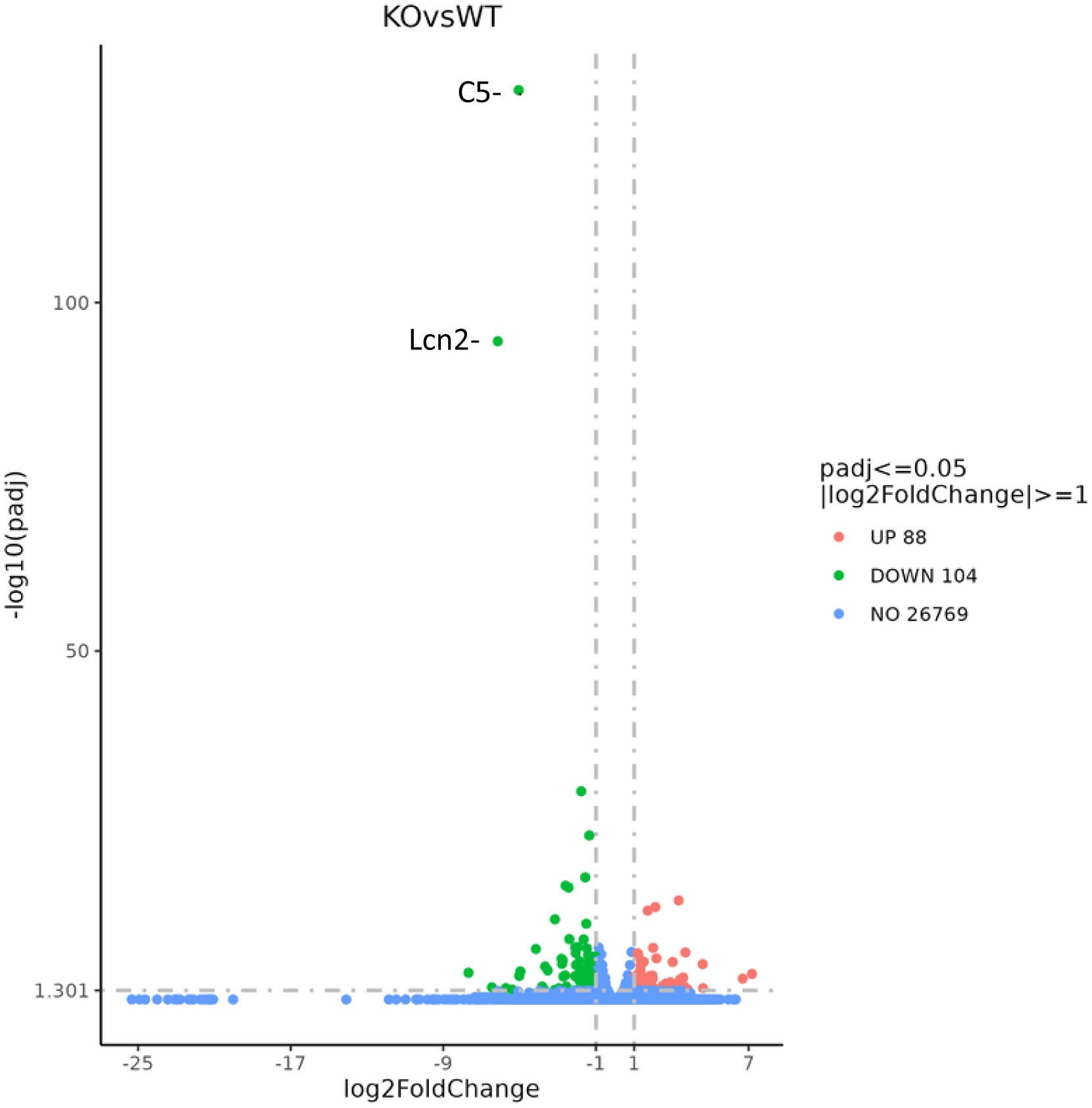
RNA-seq analysis of lung tissue from crTMPRSS2 and WT rats. Volcano plot showing differential gene expression in lung tissue of crTMPRSS2 rats compared with wild-type (WT) controls. Significantly upregulated genes are shown in red and significantly downregulated genes in green (padj ≤ 0.05, |log2FoldChange| ≥ 1), while non-significant genes are shown in blue. Among the downregulated genes were inflammatory and immune-related markers including C5 and Lcn2.

**Figure 11.**
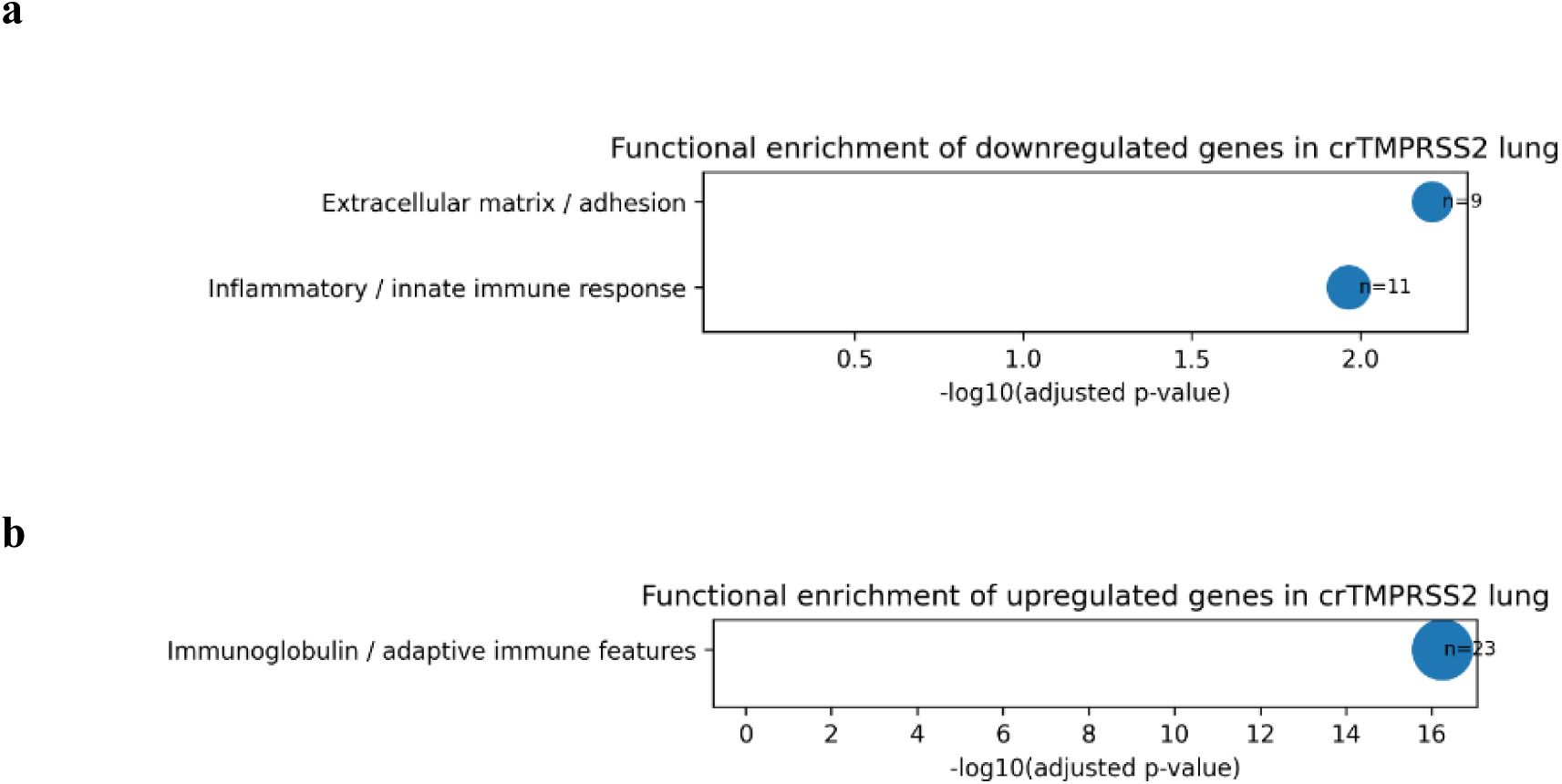
Functional enrichment of differentially expressed genes in crTMPRSS2 lung. **(a)** Enrichment analysis based on gene annotation and functional grouping of downregulated genes in crTMPRSS2 lung tissue, showing major affected categories in extracellular matrix/adhesion and inflammatory response. **(b)** Enrichment analysis of upregulated genes, highlighting immunoglobulin/adaptive immune features. Bubble size indicates gene count, and the x-axis shows −log10(adjusted p-value).

**Figure 12.**
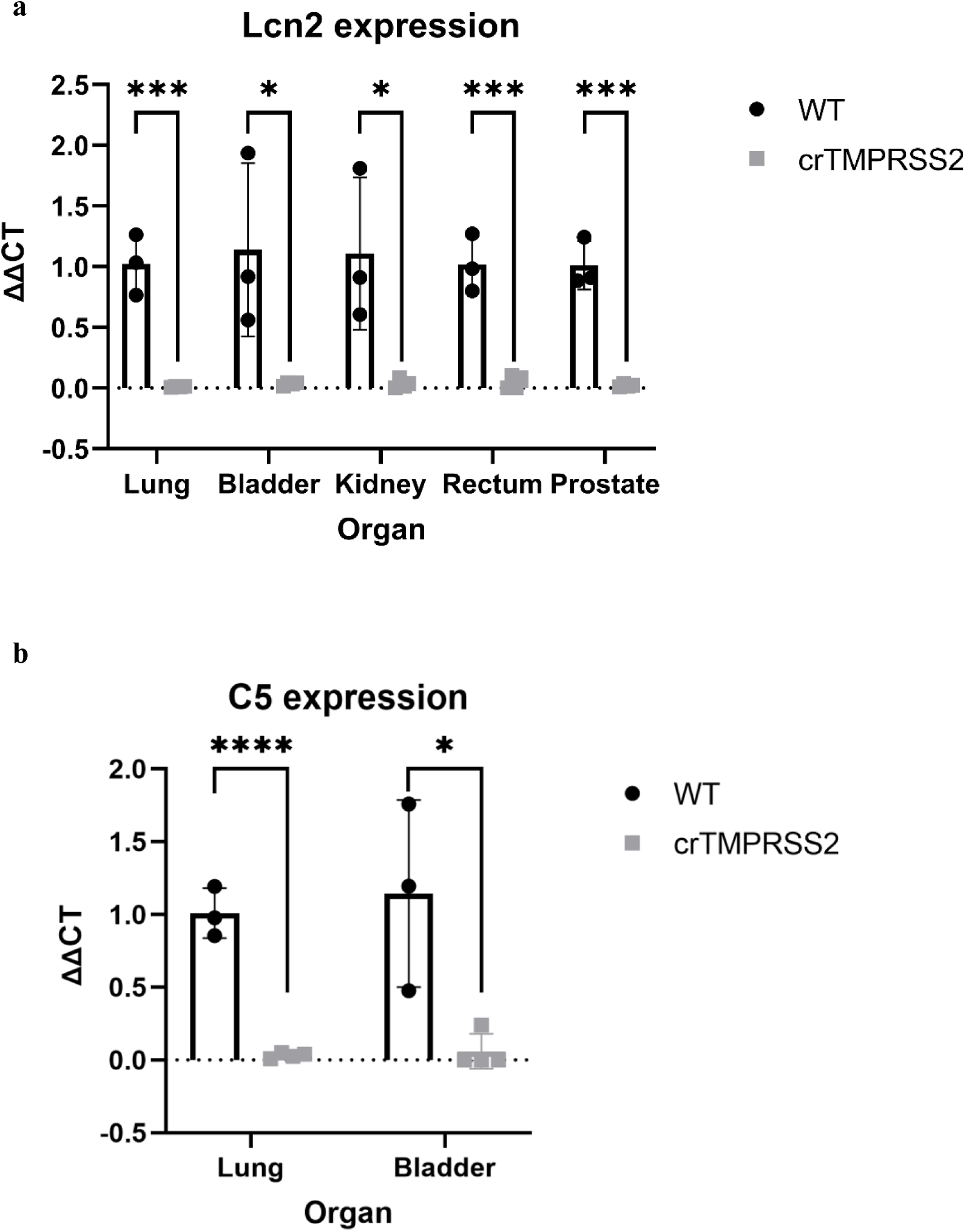
Reduced expression of C5 and Lcn2 in crTMPRSS2 rats. (a) RT-qPCR analysis of Lcn2 expression in lung, bladder, kidney, rectum, and prostate of WT and crTMPRSS2 rats. Lcn2 mRNA levels were significantly decreased in all tissues analyzed, including lung (p = 0.000398), bladder (p = 0.0239), kidney (p = 0.0165), rectum (p = 0.000446), and prostate (p = 0.000154). (b) RT-qPCR analysis of C5 expression in lung and bladder of WT and crTMPRSS2 rats. C5 mRNA levels were significantly reduced in crTMPRSS2 animals compared with WT controls in lung (p = 7.80 × 10⁻⁵) and bladder (p = 0.0193).

**Figure 13.**
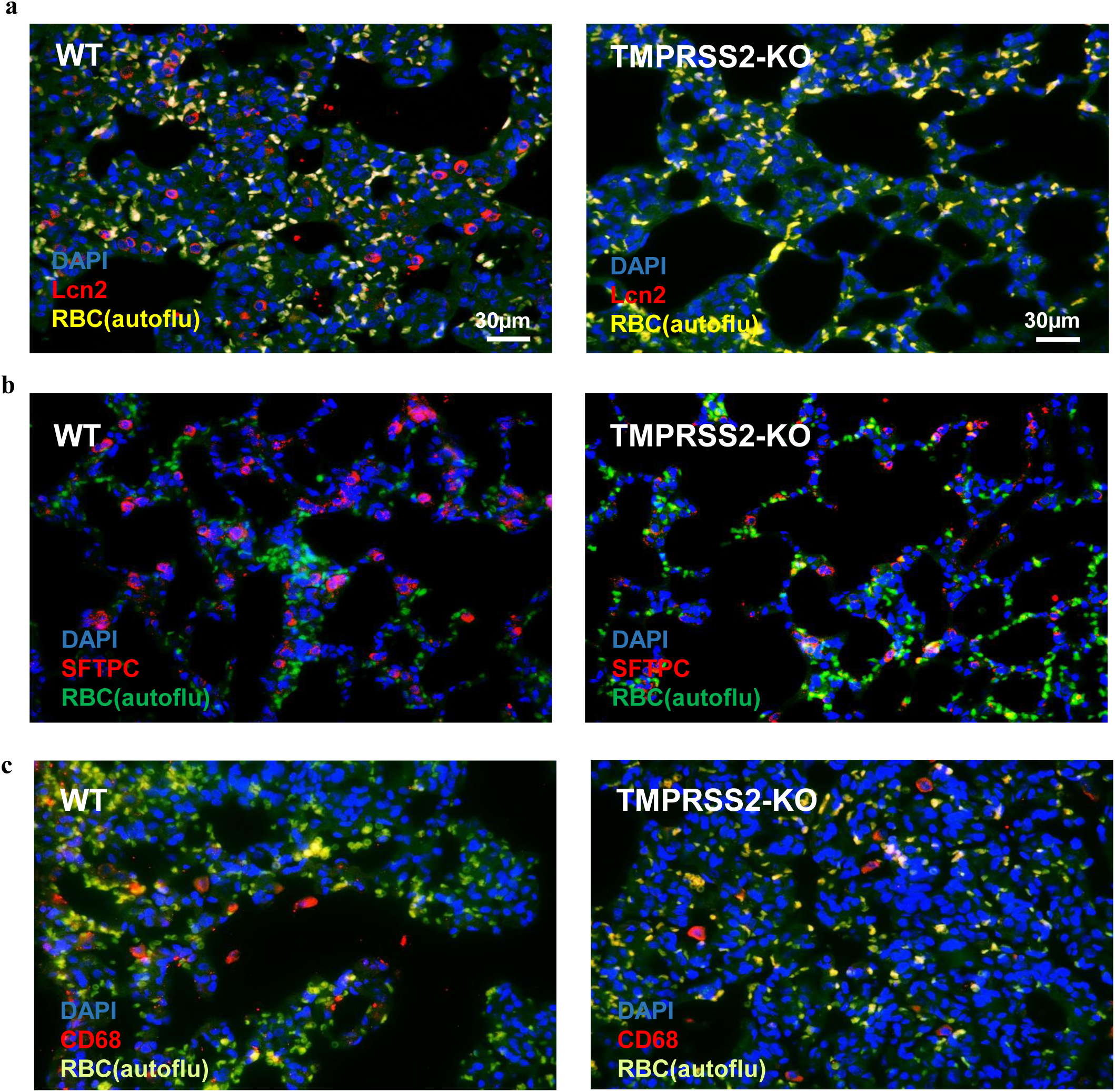
Immunofluorescence staining of inflammatory and epithelial markers in WT and TMPRSS2-KO lung tissue. Representative immunofluorescence images of lung sections from WT and TMPRSS2-KO animals stained for Lcn2/NGAL (a), SFTPC (b), and CD68 (c). Nuclei were counterstained with DAPI. Red blood cell autofluorescence is shown as indicated. Lcn2 staining was strongly reduced in TMPRSS2-KO lungs compared with WT controls. CD68-positive cells were detected in both genotypes but appeared reduced in alveolar regions of TMPRSS2-KO lungs. SFTPC staining was present in both WT and TMPRSS2-KO lung tissue. Scale bars: 30 µm. n = 3 animals per group, with 2–3 sections analyzed per animal.

## Discussion

TMPRSS2 is an important host protease in SARS-CoV-2 infection, classically viewed as a cell-surface factor that primes the spike protein for membrane fusion. Our results extend this view by defining mechanistic features of TMPRSS2 maturation and by showing that activated TMPRSS2 directly shapes ACE2 processing, trafficking, and release.

In this study, we found that TMPRSS2 autoactivation can occur in trans. Co-expression of active TMPRSS2 with an enzymatically inactive TMPRSS2i led to cleavage of TMPRSS2i, detected as release of the FLAG-tagged serine protease domain confirning a recent study^31^. This intermolecular activation suggests that TMPRSS2 may assemble into dimers or higher-order complexes during maturation. A similar principle has been reported for the related airway transmembrane serine protease TMPRSS11A, which undergoes autoactivation in-trans, supporting the idea that intermolecular activation may be a shared feature within the TTSP family^31,37^.

Our data also showed that binding to ACE2 is linked to the activation state of TMPRSS2. Only catalytically active TMPRSS2 co-precipitated with ACE2 confirming a previous report^4^ and, when interacting with TMPRSS2, ACE2 underwent two changes: an ∼85 kDa N-terminal fragment consistent with cleavage and an ∼5 kDa mobility shift. The ∼5 kDa mobility shift has also been observed in earlier studies coexpressing ACE2 and TMPRSS2, but was never further investigated^5,10^. Both changes required TMPRSS2 protease activity, and importantly, del1–83TMPRSS2 separated the two phenotypes by inducing the mobility shift but much less the cleavage. This argues that ACE2 cleavage and the mobility shift arise from separable mechanisms and that the cytoplasmic TMPRSS2 N-terminus contributes specifically to productive ACE2 proteolysis, likely by controlling localization of TMPRSS2. There are no studies yet on the function of the N-terminal domain of TMPRSS2 but previous studies suggest that intracellular regions of the TTSP family of enzymes contribute to their protein maturation, intracellular trafficking, and membrane localization^38^.

TMPRSS2 markedly altered ACE2 subcellular distribution and surface availability as also described earlier^4^. ACE2-GFP expressed alone localized prominently to the plasma membrane, whereas co-expression with active TMPRSS2 caused loss of clear membrane localization and an intracellular pattern that correlated with ER staining. Membrane biotinylation supported this observation, showing that surface ACE2 was strongly reduced in the presence of active TMPRSS2, while ACE2 remained detectable at the membrane when expressed alone or with TMPRSS2i. In parallel, ACE2 release into the culture medium increased upon co-expression with TMPRSS2, consistent with TMPRSS2-dependent shedding contributing to depletion of membrane ACE2. In conclusion, the expression level of TMPRSS2 in a cell determines the concentration of active ACE2 on its surface and the extent of its shedding.

We propose a working model in which full-length ACE2, normally trafficked through the ER/Golgi to the plasma membrane, encounters activated TMPRSS2 early in the secretory pathway, before full post-translational maturation. In this context, TMPRSS2 association could promote ACE2 cleavage within the collectrin-like domain to generate soluble ACE2 that is subsequently released, explaining the reduction of surface ACE2 and the appearance of ACE2 in the medium.

A substantial body of prior research has demonstrated the critical role of N-glycosylation in regulating ACE2 cell surface expression. Furthermore, studies have demonstrated that N-glycosylation deficiency can induce the accumulation of ACE2 within the ER^39^. Therefore, we speculated that TMPRSS2 binding structurally hinders glycan addition or glycan maturation on ACE2. This would be, consistent with the reduction of the ACE2 surface localization in the fully deglycosylated ACE2 (N1–N7Q) which resembled WT ACE2 in the presence of TMPRSS2, supporting the idea that global glycan loss can phenocopy a TMPRSS2-bound state with impaired surface trafficking.

These conclusions are based on transient overexpression in cultured cell lines, which may accentuate retention or processing, and co-precipitation does not by itself distinguish direct binding from association within larger complexes. Future experiments testing glycan maturation directly (e.g., Endo H/PNGase F sensitivity and glycoproteomics), titrating TMPRSS2 expression, quantifying ACE2 surface levels by flow cytometry, or probing TMPRSS2 dimerization or oligomerization will refine the proposed mechanism. Overall, our results support that TMPRSS2 can autoactivate in trans and, once activated, associates with ACE2 to cleave it and remodel ACE2 processing, leading to reduced ACE2 surface compartmentalization and increased ACE2 release.

The second part of our study assessed the *in-vivo* relevance of the cell culture findings. To this purpose we generated a TMPRSS2-deficient rat model using CRISPR/Cas9 mediated gene editing which introduced a frame shift mutation into exon 2 of the *Tmprss2* gene. The absence of the protein was evidenced by a marked reduction in total TTSP protease activity in several organs and a failure to detect any peptide of TMPRSS2 in the proteomics analysis in TMPRSS2-KO rat lungs while it was clearly detectable in all WT samples. In concordance with our cell culture findings revealing TMPRSS2 as sheddase for ACE2, ACE2 activity was increased in tissues and decreased in the blood of the KO rats. It is well known that ACE2 is released from tissue in inflammatory conditions. Several studies have implicated ADAM17 and ADAM10, both belonging to the ADAM (a disintegrin and metalloproteinase) family, in this shedding process^40–44^. Our study adds TMPRSS2 as an additional candidate sheddase which is at least responsible to release a part of ACE2 constitutively from tissues into the blood. Its role in induced shedding in pathological processes needs future experimental clarification. However, such a role is supported by its reported upregulation in inflammation and diabetes^45–47^.

Concordant with an anti-inflammatory role of ACE2 in the lung, TMPRSS2-deficient rats showed a clear reduction in the baseline expression of inflammation markers. In particular Lcn2, one of the most relevant inflammation markers in the lung, was drastically reduced on the mRNA and protein level. We don’t know yet whether this is, indeed, the consequence of the preserved tissue localization of ACE2 or an independent effect exerted by the absence of TMPRSS2 in this organ. At least, it has been repeatedly shown that ACE2 inhibits lung and kidney inflammation and suppresses Lcn2 expression^48–55^. Future studies are warranted to confirm that TMPRSS2 inhibition or deletion protects lung and kidney from injury by preserving the tissue localization of ACE2.

## References

1. Huang, C. et al. Clinical features of patients infected with 2019 novel coronavirus in Wuhan, China. The Lancet 395, 497–506 (2020).

2. Wu, F. et al. A new coronavirus associated with human respiratory disease in China. Nature 579, 265–269 (2020).

3. WHO Coronavirus (COVID-19) Dashboard. https://covid19.who.int.

4. Shulla, A. et al. A Transmembrane Serine Protease Is Linked to the Severe Acute Respiratory Syndrome Coronavirus Receptor and Activates Virus Entry. J. Virol. 85, 873–882 (2011).

5. Heurich, A. et al. TMPRSS2 and ADAM17 Cleave ACE2 Differentially and Only Proteolysis by TMPRSS2 Augments Entry Driven by the Severe Acute Respiratory Syndrome Coronavirus Spike Protein. J. Virol. 88, 1293–1307 (2014).

6. Li, W. et al. Angiotensin-converting enzyme 2 is a functional receptor for the SARS coronavirus. Nature 426, 450–454 (2003).

7. Bader, M. ACE2, angiotensin-(1–7), and Mas: the other side of the coin. Pflüg. Arch. - Eur. J. Physiol. 465, 79–85 (2013).

8. Santos, R. A. S. et al. The ACE2/Angiotensin-(1–7)/MAS Axis of the Renin-Angiotensin System: Focus on Angiotensin-(1–7). Physiol. Rev. 98, 505–553 (2018).

9. McCallum, M. et al. TMPRSS2-mediated coronavirus spike activation and inhibition. Nat. Struct. Mol. Biol. 33, 810–823 (2026).

10. Essalmani, R. et al. Distinctive Roles of Furin and TMPRSS2 in SARS-CoV-2 Infectivity. J. Virol. 96, e00128–22 (2022).

11. Hoffmann, M. et al. SARS-CoV-2 Cell Entry Depends on ACE2 and TMPRSS2 and Is Blocked by a Clinically Proven Protease Inhibitor. Cell 181, 271–280.e8 (2020).

12. Zhang, L., Hoffmann, M. & Pöhlmann, S. Lock out: targeting TMPRSS2 to block influenza and coronaviruses. J. Virol. e0080725 (2026) doi:10.1128/jvi.00807-25.

13. Bugge, T. H., Antalis, T. M. & Wu, Q. Type II Transmembrane Serine Proteases. J. Biol. Chem. 284, 23177–23181 (2009).

14. Afar, D. E. H. et al. Catalytic Cleavage of the Androgen-regulated TMPRSS2 Protease Results in Its Secretion by Prostate and Prostate Cancer Epithelia. Cancer Res. 61, 1686–1692 (2001).

15. Saunders, N. et al. TMPRSS2 is a functional receptor for human coronavirus HKU1. Nature 624, 207–214 (2023).

16. Sakai, K. et al. The Host Protease TMPRSS2 Plays a Major Role in In Vivo Replication of Emerging H7N9 and Seasonal Influenza Viruses. J. Virol. 88, 5608 (2014).

17. Tarnow, C. et al. TMPRSS2 Is a Host Factor That Is Essential for Pneumotropism and Pathogenicity of H7N9 Influenza A Virus in Mice. J. Virol. 88, 4744–4751 (2014).

18. Heindl, M. R. et al. ACE2 acts as a novel regulator of TMPRSS2-catalyzed proteolytic activation of influenza A virus in airway cells. J. Virol. 98, e00102 (2024).

19. Ciacci Zanella, G., et al. Pigs lacking TMPRSS2 displayed fewer lung lesions and reduced inflammatory response when infected with influenza A virus. Front. Genome Ed. 5, 1320180 (2023).

20. Lambertz, R. L. O. et al. H2 influenza A virus is not pathogenic in Tmprss2 knock-out mice. Virol. J. 17, 56 (2020).

21. Lambertz, R. L. O. et al. Tmprss2 knock-out mice are resistant to H10 influenza A virus pathogenesis. J. Gen. Virol. 100, 1073–1078 (2019).

22. Hatesuer, B. et al. Tmprss2 is essential for influenza H1N1 virus pathogenesis in mice. PLoS Pathog. 9, e1003774 (2013).

23. Iwata-Yoshikawa, N. et al. Essential role of TMPRSS2 in SARS-CoV-2 infection in murine airways. Nat. Commun. 13, 6100 (2022).

24. Iwata-Yoshikawa, N. et al. TMPRSS2 Contributes to Virus Spread and Immunopathology in the Airways of Murine Models after Coronavirus Infection. J. Virol. 93, e01815–18 (2019).

25. Shuai, H. et al. An orally available Mpro/TMPRSS2 bispecific inhibitor with potent anti-coronavirus efficacy in vivo. Nat. Commun. 16, 6541 (2025).

26. Barros de Lima, G., et al. TMPRSS2 as a Key Player in Viral Pathogenesis: Influenza and Coronaviruses. Biomolecules 15, 75 (2025).

27. Zhang, Y. et al. Transmembrane serine protease TMPRSS2 implicated in SARS-CoV-2 infection is autoactivated intracellularly and requires N-glycosylation for regulation. J. Biol. Chem. 298, 102643 (2022).

28. Popova, E., Krivokharchenko, A., Ganten, D. & Bader, M. Efficiency of transgenic rat production is independent of transgene-construct and overnight embryo culture. Theriogenology 61, 1441–1453 (2004).

29. Demichev, V., Messner, C. B., Vernardis, S. I., Lilley, K. S. & Ralser, M. DIA-NN: neural networks and interference correction enable deep proteome coverage in high throughput. Nat. Methods 17, 41–44 (2020).

30. Ritchie, M. E. et al. limma powers differential expression analyses for RNA-sequencing and microarray studies. Nucleic Acids Res. 43, e47 (2015).

31. Végh, B. et al. Efficient production of fully active, SARS-CoV-2-priming, wildtype TMPRSS2 ectodomain via co-expression of HAI-2 allows for both auto- and cross-activation mechanisms. Biochem. J. 482, 1993–2010 (2025).

32. Paoloni-Giacobino, A., Chen, H., Peitsch, M. C., Rossier, C. & Antonarakis, S. E. Cloning of the TMPRSS2 Gene, Which Encodes a Novel Serine Protease with Transmembrane, LDLRA, and SRCR Domains and Maps to 21q22.3. Genomics 44, 309–320 (1997).

33. Supekar, N. T. et al. Variable posttranslational modifications of severe acute respiratory syndrome coronavirus 2 nucleocapsid protein. Glycobiology 31, 1080–1092 (2021).

34. Tipnis, S. R. et al. A Human Homolog of Angiotensin-converting Enzyme: Cloning and functional expression as a captopril-insensitive carboxypeptidase. J. Biol. Chem. 275, 33238–33243 (2000).

35. Shajahan, A. et al. Comprehensive characterization of N- and O-glycosylation of SARS-CoV-2 human receptor angiotensin converting enzyme 2. Glycobiology cwaa101 (2020) doi:10.1093/glycob/cwaa101.

36. Isobe, A. et al. ACE2 N-glycosylation modulates interactions with SARS-CoV-2 spike protein in a site-specific manner. Commun. Biol. 5, 1188 (2022).

37. Zhang, C. et al. Intracellular autoactivation of TMPRSS11A, an airway epithelial transmembrane serine protease. J. Biol. Chem. 295, 12686–12696 (2020).

38. Hooper, J. D., Clements, J. A., Quigley, J. P. & Antalis, T. M. Type II Transmembrane Serine Proteases: Insights into an emerging class of cell surface proteolytic enzymes. J. Biol. Chem. 276, 857–860 (2001).

39. Rowland, R. & Brandariz-Nuñez, A. Analysis of the Role of N-Linked Glycosylation in Cell Surface Expression, Function, and Binding Properties of SARS-CoV-2 Receptor ACE2. Microbiol. Spectr. 9, e01199-21 (2021).

40. Lambert, D. W. et al. Tumor necrosis factor-alpha convertase (ADAM17) mediates regulated ectodomain shedding of the severe-acute respiratory syndrome-coronavirus (SARS-CoV) receptor, angiotensin-converting enzyme-2 (ACE2). J. Biol. Chem. 280, 30113–30119 (2005).

41. Webers, M. et al. The metalloproteinase ADAM10 sheds angiotensin-converting enzyme (ACE) from the pulmonary endothelium as a soluble, functionally active convertase. FASEB J. 38, e70105 (2024).

42. Patel, V. B. et al. Angiotensin II induced proteolytic cleavage of myocardial ACE2 is mediated by TACE/ADAM-17: a positive feedback mechanism in the RAS. J. Mol. Cell. Cardiol. 66, 167–176 (2014).

43. Xu, J. et al. Clinical Relevance and Role of Neuronal AT1 Receptors in ADAM17-Mediated ACE2 Shedding in Neurogenic Hypertension. Circ. Res. 121, 43–55 (2017).

44. Niehues, R. V. et al. The collectrin-like part of the SARS-CoV-1 and −2 receptor ACE2 is shed by the metalloproteinases ADAM10 and ADAM17. FASEB J. 36, e22234 (2022).

45. Batchu, S. N. et al. Lung and Kidney ACE2 and TMPRSS2 in Renin-Angiotensin System Blocker-Treated Comorbid Diabetic Mice Mimicking Host Factors That Have Been Linked to Severe COVID-19. Diabetes 70, 759–771 (2021).

46. Nowak, J. K. et al. Age, Inflammation, and Disease Location Are Critical Determinants of Intestinal Expression of SARS-CoV-2 Receptor ACE2 and TMPRSS2 in Inflammatory Bowel Disease. Gastroenterology 159, 1151–1154.e2 (2020).

47. Visvanathan, R., Houghton, M. J. & Williamson, G. Impact of glucose, inflammation and phytochemicals on ACE2, TMPRSS2 and glucose transporter gene expression in human intestinal cells. Antioxidants 14, 253 (2025).

48. Sanad, A. M. et al. Transgenic angiotensin-converting enzyme 2 overexpression in the rat vasculature protects kidneys from ageing-induced injury. Kidney Int. 104, 293–304 (2023).

49. Imai, Y. et al. Angiotensin-converting enzyme 2 protects from severe acute lung failure. Nature 436, 112–116 (2005).

50. Bae, E. H. et al. Murine recombinant angiotensin-converting enzyme 2 attenuates kidney injury in experimental Alport syndrome. Kidney Int. 91, 1347–1361 (2017).

51. Yamaguchi, T. et al. ACE2-like carboxypeptidase B38-CAP protects from SARS-CoV-2-induced lung injury. Nat. Commun. 12, 6791 (2021).

52. Shirazi, M. et al. Altered kidney distribution and loss of ACE2 into the urine in acute kidney injury. Am. J. Physiol. Renal Physiol. 327, F412–F425 (2024).

53. Rey-Parra, G. J. et al. Angiotensin converting enzyme 2 abrogates bleomycin-induced lung injury. J. Mol. Med. 90, 637–647 (2012).

54. Gu, H. et al. Angiotensin-converting enzyme 2 inhibits lung injury induced by respiratory syncytial virus. Sci. Rep. 6, 19840 (2016).

55. Hassler, L. et al. A Novel Soluble ACE2 Protein Provides Lung and Kidney Protection in Mice Susceptible to Lethal SARS-CoV-2 Infection. J. Am. Soc. Nephrol. 33, 1293–1307 (2022).

